# Transcription-dependent spreading of canonical yeast GATA factor across the body of highly expressed genes

**DOI:** 10.1101/238550

**Authors:** Aria Ronsmans, Maxime Wery, Camille Gautier, Marc Descrimes, Evelyne Dubois, Antonin Morillon, Isabelle Georis

**Author notes:** These authors contributed equally to this work (Aria Ronsmans, Maxime Wery). Corresponding authors (IG), (AM).

## Abstract

GATA transcription factors are highly conserved among eukaryotes and play roles in transcription of genes implicated in cancer progression and hematopoiesis. However, although their consensus binding sites have been well defined *in vitro*, the *in vivo* selectivity for recognition by GATA factors remains poorly characterized. Using ChIP-Seq, we identified the Dal80 GATA factor targets in yeast. Our data reveal Dal80 binding to a large set of promoters, sometimes independently of GATA sites. Strikingly, Dal80 was also detected across the body of promoter-bound genes, correlating with high, Dal80-sensitive expression. Mechanistic single-gene experiments showed that Dal80 spreading across gene bodies is independent of intragenic GATA sites but requires transcription elongation. Consistently, Dal80 co-purified with the post-initiation form of RNA Polymerase II. Our work suggests that GATA factors could play dual, synergistic roles during transcription initiation and post-initiation steps, promoting efficient remodeling of the gene expression program in response to environmental changes.

**Author Summary:** GATA transcription factors are highly conserved among eukaryotes and play key roles in cancer progression and hematopoiesis. In budding yeast, four GATA transcription factors are involved in the response to the quality of nitrogen supply. We have determined the whole genome binding profile of one of them, Dal80, and revealed that it also binds across the body or promoter-bound genes. Our observation that ORF binding correlated with elevated transcription levels and exquisite Dal80 sensitivity suggests that GATA factors could play other, unexpected roles at post-initiation stages in eukaryotes.

## Introduction

In eukaryotes, gene transcription by RNA polymerase II (PolII) is initiated by the binding of specific transcription factors to double stranded DNA. The yeast transcription factors target regulatory regions called UAS or URS (for *U*pstream *A*ctivating/*R*epressing *S*equences), generally directly adjacent to the core promoter. The generated regulatory signals converge at the core promoter where they permit the regulation of PolII recruitment via the ‘TATA box-binding protein’ and associated general transcription factors [1,2]. The transcription factor binding sites are usually short sequences ranging from 8 to 20 bp [3]. They are most often similar but generally not identical, differing by some nucleotides from one another [3], making it sometimes difficult to predict whether a given UAS will function as such *in vivo*.

GATA factors constitute a family of transcription factors highly conserved among eukaryotes and characterized by the presence of a DNA binding domain which consists of four cysteines (fitting the consensus sequence CX_2_CX_17-18_CX_2_C) coordinating a zinc ion followed by a basic carboxy-terminal tail [4]. *In vitro*, this conserved binding domain allows the recognition of consensus GATA DNA sequences (GATAA, GATAAG or GATTAG), found in the promoter of numerous genes, which was demonstrated in various organisms either with direct or indirect methods [4–8]. Structure elucidation of GATA factor binding domains in complex with their DNA target suggest an important role for the basic tail in sequence specificity determination [4,8].

Since its discovery 40 years ago in chicken cells, the family of GATA factors was extended in human cells and represents master regulators of hematopoiesis and cancer [9]. However, although approximately 7 million GATA motifs can be found in the human genome, the GATA factors occupy only 0.1–1% of them. Conversely, other regions are occupied by GATA factors despite lacking the consensus motif [10,11]. In addition, GATA factors can swap among them for the same motif and switch from active or repressive transcriptional activity. All these observations raised the main paradigm on understanding the selective recruitment and activity of GATA factors and what the determinants of their residency on the chromatin are [12,13].

In yeast, the family of GATA transcription factors contains over 10 members [14]. Four of them are implicated in the regulation of *N*itrogen *C*atabolite *R*epression- (NCR-) sensitive genes, the expression of which is repressed in the presence of a preferred nitrogen source (glutamine, asparagine, ammonia) and derepressed when only poor nitrogen sources (e.g. proline, leucine, urea) are available [15]. These key GATA factors involved in NCR signaling are two activators (Gln3 and Gat1/Nil1) and two repressors (Gzf3/Nil2/Deh1 and Dal80/Uga43) [16–21]. Moreover, the expression of *DAL80* and *GAT1* is also NCR-sensitive, which implies cross- and autogenous regulations of the GATA factors in the NCR mechanisms [21–24]. Under nitrogen limitation, expression of *DAL80* is highly induced [18], and Dal80 enters the nucleus where it competes with the two GATA activators for the same binding sites [5,22,25]. Although initially described as being active under nitrogen abundance [20,21], the Gzf3 repressor also localizes to NCR-sensitive promoters in conditions of activation [23].

*In vitro*, the Gln3 and Gat1 activators bind, presumably as monomers, to single GATA sequences [26], like their orthologous vertebrate counterparts. On the other hand, the Dal80 repressor can form homo-and heterodimers with Gzf3, thanks to the presence of an N-terminal leucine zipper [27] and, consistently, Dal80 binds *in vitro* to two GATA sequences [5,22,27,28]. *In vivo*, GATA factor binding site recognition also appears to require repeats of GATA motifs within promoters. Indeed, synthetic constructions containing a portion of the NCR-sensitive *DAL5* promoter have shown in a β-galactosidase assay that at least two copies of the consensus are required to support efficient gene activation. This led to the actual fuzzy definition of *UAS_NTR_*, consisting in two GATA sites located sufficiently close to one another to present a binding platform for GATA factors [29–31].

*In vivo*, GATA ChIP analyses have been carried out genome-wide in yeast [32–35]. They allowed gaining insights into the GATA factor gene network through the identification of direct targets and characterization of the DNA elements required to render a gene NCR-sensitive. However, these studies were not performed in activating conditions, when all GATA factors are expressed, localized in the nucleus and active, so that the current list of GATA factor targets are likely to be underestimated. On another hand, bioinformatic analyses have shown that, since GATA sequences are short, they can be found almost everywhere throughout the genome. Therefore, based on the sole criteria of the presence of repeated GATA sequences in yeast promoters, a third of the yeast genes could hypothetically be NCR regulator targets [36]. Such GATA motif repetitions have been found in the promoter of 91 genes, inducible by GATA activators in absence of a good nitrogen source, supposed to be directly targeted by the GATA activators [37]. Nevertheless, the functionality of these supposed UAS still needs to be directly demonstrated *in vivo* [1].

Here, we provide the first genome-wide identification of Dal80 targets in yeast, in physiological conditions where Dal80 is fully expressed and active. Using a ChIP-Seq approach combined to a bioinformatic peak-calling procedure, we defined the exhaustive set of Dal80-bound promoters, which turned out to be much larger than anticipated. Our data indicate that at some promoters, Dal80 recruitment occurs independently of GATA sites. Strikingly, Dal80 was also detected across the body of a subset of genes bound at the promoter, globally correlating with high and Dal80-sensitive expression. Mechanistic single-gene experiments confirmed the Dal80 binding profiles, further indicating that Dal80 spreading across gene bodies occurs independently of intragenic GATA sites but requires transcription elongation. Finally, co-immunoprecipitation experiments revealed that Dal80 physically interacts with both intitiating and elongating form of PolII. Our work suggests that in addition to their well-established role at the (pre)inititation stage, GATA factors could be involved in transcription elongation, promoting efficient remodeling of the gene expression program in response to environmental changes.

## Results

### Genome-wide identification of Dal80-bound promoters

In order to determine the genome-wide occupancy of a GATA factor in yeast, our rationale was to choose Dal80 as it is known to be highly expressed in derepressing conditions and forms chromosome foci when tagged by GFP [38]. We thus grew yeast cells in proline-containing media and performed a ChIP-Seq analysis using a Dal80-Myc^13^ tagged strain and the isogenic untagged strain, as a control (Fig 1A). Dal80-bound regions were identified using a peak-calling algorithm (see Material & Methods). A promoter was defined as bound by Dal80 when >75% of the −350 to −100 region (relative to the downstream ORF start site) was overlapped by a peak (Fig 1B). We chose to use as the reference coordinate the translation initiation codon rather than the transcription start site (TSS) because the latter has not been accurately determined for all genes. Then, our arbitrary definition of the promoter as the −350 to −100 region relative to the ATG codon was based on the distribution of the TSS-ATG distance for genes with an annotated TSS (median and average distance = 58 and 107 bp, respectively; see S1 Fig).

**Fig 1.**
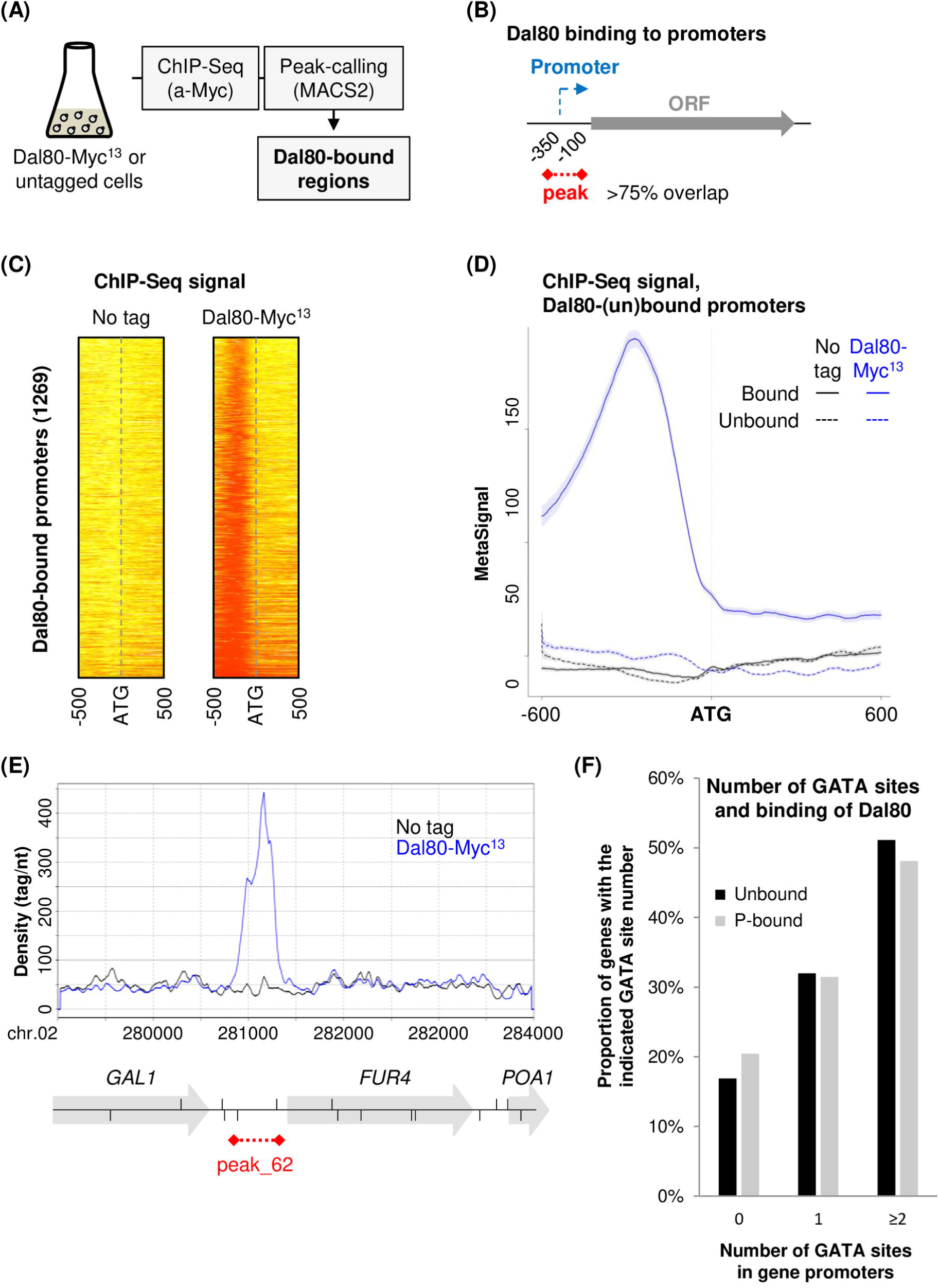
Genome-wide identification of Dal80-bound promoters. (A) Overview of the ChIP-Seq analysis. Biological duplicates of FV78 (Dal80-Myc^13^) and 25T0b (no tag) cells were grown to mid-log phase in proline-containing medium, and then harvested. After chromatin extraction and sonication, Dal80-Myc^13^ was immunoprecipitated using α-Myc antibody. Co-precipitated DNA fragments were purified and used to construct ChIP-Seq libraries. After sequencing of the libraries, signals were computed using uniquely mapped reads. Dal80-bound regions were identified using a peak-calling procedure using MACS2. (B) Identification of Dal80-bound promoters. After peak-calling, Dal80-bound promoters were identified on the basis of a >75% overlap of the −100 to −350 regions (relative to the downstream ORF start site) by the peak (represented in as a red dashed line). (C) Heatmap view of the ChIP-Seq signal in the ATG +/− 500 bp region for the 1269 genes identified as bound by Dal80 at the promoter, in untagged (left) and Dal80-Myc^13^ cells (right). The colour goes from yellow to red as the signal increases. (D) Metagene view of the ChIP-Seq signal along the ATG +/- 600 bp region for the 1269 genes identified as bound by Dal80 at the promoter (solid lines) and for the unbound genes (dashed lines), in untagged (black) and Dal80-Myc^13^ cells (blue). For each group of genes, normalized coverage (tag/nt) for each gene was piled up, and the average signal was plotted. The shading surrounding each line denotes the 95% confidence interval. (E) Snapshot of ChIP-Seq signals along the *FUR4 locus*. Densities (tag/nt) are shown for the untagged (black line) and Dal80-Myc^13^ (blue line) strains. ORFs and the Dal80-Myc^13^ ChIP-Seq peak are represented by grey arrows and the red dashed line, respectively. The position (and orientation) of each GATA site is represented by vertical segments above (sense GATA sites) or below (antisense GATA sites) the *locus* line. The snapshot was produced using the VING software (Descrimes *et al.*, 2015). (F) Number of GATA sites in the promoter of Dal80-unbound (4522) and promoter-bound (1268) genes. The analysis was performed using RSAT (Medina-Rivera *et al.*, 2015), across the −500 to −1 region (relative to the ATG codon of the downstream ORF).

Strikingly, Dal80 was found to bind to 1269 gene promoters (Fig 1C–D and S1 Table). Fig 1E shows a snapshot of ChIP-Seq signals at the promoter of the *FUR4* gene, identified as bound by Dal80. The number of Dal80-bound promoters, corresponding to 22% of all protein-coding genes promoters, is much higher than expected given the hundred target genes that are generally cited for the GATA transcriptional activators [37,39], that are presumed to bind to the same sites as Dal80. However, we noted that some peaks (221) overlapped with several promoters (471), mainly of divergent genes (442). In such cases, it is therefore possible that only one of the two divergent promoters is indeed targeted by Dal80, so that the number of Dal80-bound promoters could actually be a little less important than 1269.

As expected, genes related to previous nitrogen regulation screens were present within our list [37,39–46](S2 Table). Among the genes showing Dal80-binding to the promoter, we also noticed a strong enrichment for cytoplasmic translation genes, as well as genes involved in small molecule biosyntheses, including amino acids (S3 Table).

Surprisingly, analysis of GATA site occurrence over Dal80-bound and unbound promoters revealed no major difference between the two classes (Fig 1F), 48.2% and 51.3% of Dal80- bound and unbound promoters containing at least two GATA sites, respectively (S4 Table). Intriguingly, 20% of Dal80-bound promoters do not contain any GATA site (Fig 1F and S4 Table), indicating that Dal80 recruitment can also occur independently of the presence of consensus GATA sites (see S1 Fig for visualization of Dal80 binding to a GATA-less promoter).

In summary, our ChIP-Seq analysis revealed that Dal80 binds to a set of promoters larger than previously expected, targeting biosynthetic functions and protein synthesis in addition to nitrogen catabolite repression.

### Dal80 binds the intragenic region of a subset of genes

Metagene analysis revealed that the genes bound by Dal80 at the promoter also display a signal along the gene body, although this intragenic signal remains globally lower than in the promoter-proximal region (Fig 1D). This observation prompted us to investigate the possibility that Dal80 also occupies the gene body, at least for a subset of genes.

We identified 189 genes showing Dal80 intragenic occupancy, according to a >75% overlap of the ORF by a Dal80-Myc^13^ peak (Fig 2A–B). Among them, 144 (76%) were also bound at the promoter (Fig 2B). Conversely, only 11% of the Dal80-bound promoters (144/1269) showed downstream intragenic binding (Fig 2B). On the other hand, 45 genes showing Dal80 intragenic binding were not bound at the promoter (Fig 2B). Hence, we distinguished four classes of genes (S5 Table): (*i*) those bound by Dal80 at the promoter only (“P” class; Fig 2C), (*ii*) those showing both promoter and intragenic binding (“P & O” class; Fig 2D), (*iii)* those bound across the ORF only (“O” class; Fig 2E), (*iv*) the unbound genes (Fig 2F). Interestingly, we noted that the global Dal80-Myc^13^ signal at the promoter was higher for the “P & O” class in comparison to the “P” class (compare Fig 2C–D).

**Fig 2.**
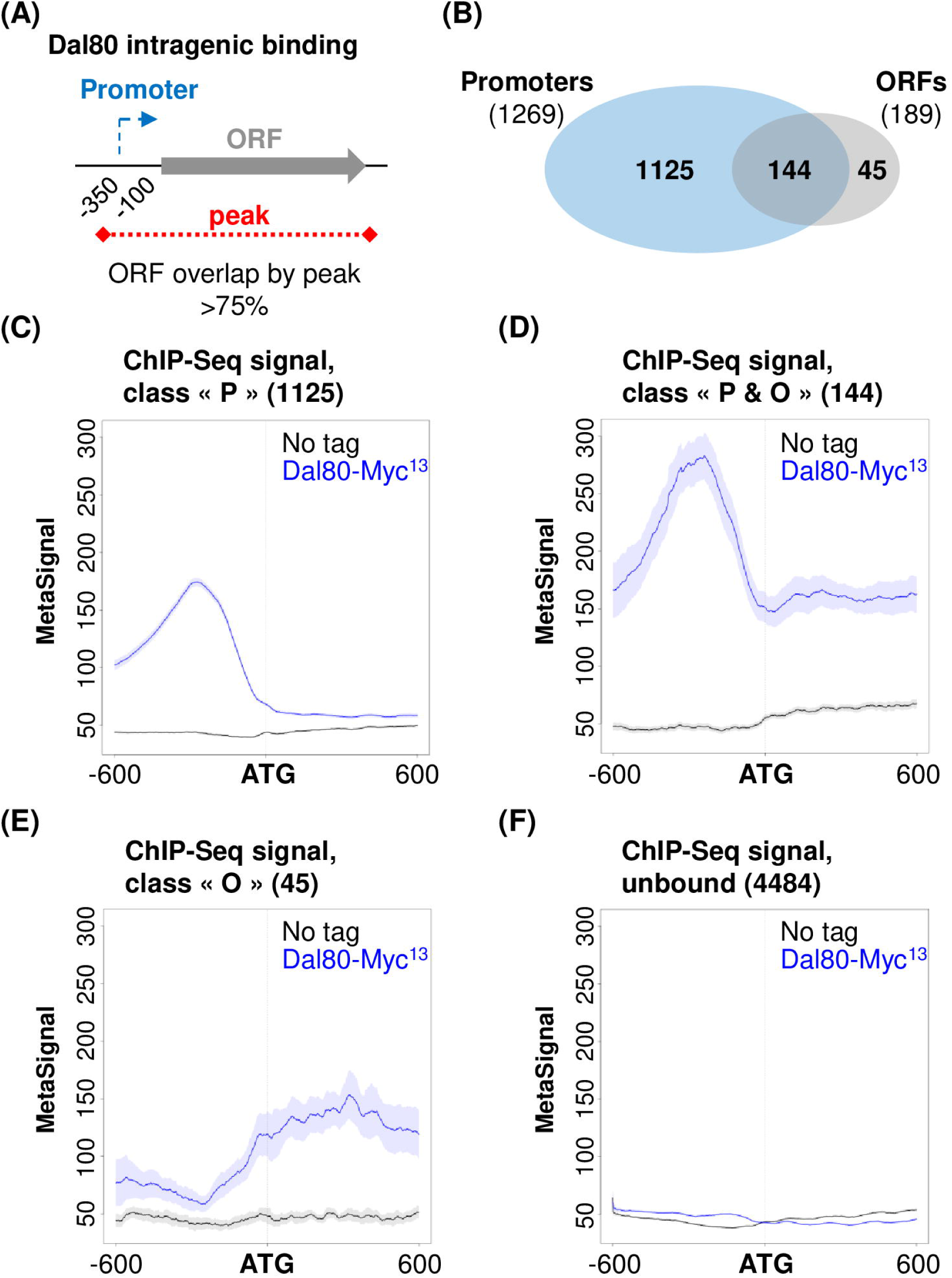
Dal80 binds the intragenic region of a subset of genes. (A) Identification of genes showing Dal80 intragenic binding. (B) Venn diagram showing the number of genes that are bound by Dal80 at the promoter and/or across the ORF. (C) Metagene view of the ChIP-Seq signal along the ATG +/- 600 bp region for the 1125 genes identified as bound by Dal80 at the promoter only (“P”), in untagged (black) and Dal80- Myc^13^ cells (blue). MetaSignal computation was as described in Fig 1d. The shading surrounding each line denotes the 95% confidence interval. (D) Same as above for the 144 genes bound by Dal80 at the promoter and across the ORF (“P & O”). (E) Same as above for the 45 genes bound by Dal80 across the ORF only (“O”). (F) Same as above for the 4484 genes that are not bound by Dal80.

In conclusion, a subset of genes displays intragenic Dal80 binding, and in most cases, this correlates with strong Dal80 recruitment at the promoter.

### Dal80 binding across gene bodies correlates with Dal80-sensitive, high expression levels

In order to correlate Dal80 binding with gene expression and Dal80-sensitivity in similar culture conditions, we performed total RNA-seq analysis in wild type and *dal80*Δ cells grown in medium with proline as the nitrogen source.

The expression of 546 genes was significantly altered in proline-grown *dal80*Δ cells compared to wild type (Fig 3A, S6 Table), among which 232 and 314 genes were negatively and positively regulated by Dal80, respectively. The first group was enriched for genes involved in small molecule catabolic processes (S7 Table), while the positively regulated genes were mostly involved in amino acid biosynthetis (S8 Table). Again, we noticed an overlap between Dal80-repressed genes and nitrogen regulated genes that were identified in other genome-wide screens (S4 Table).

**Fig 3.**
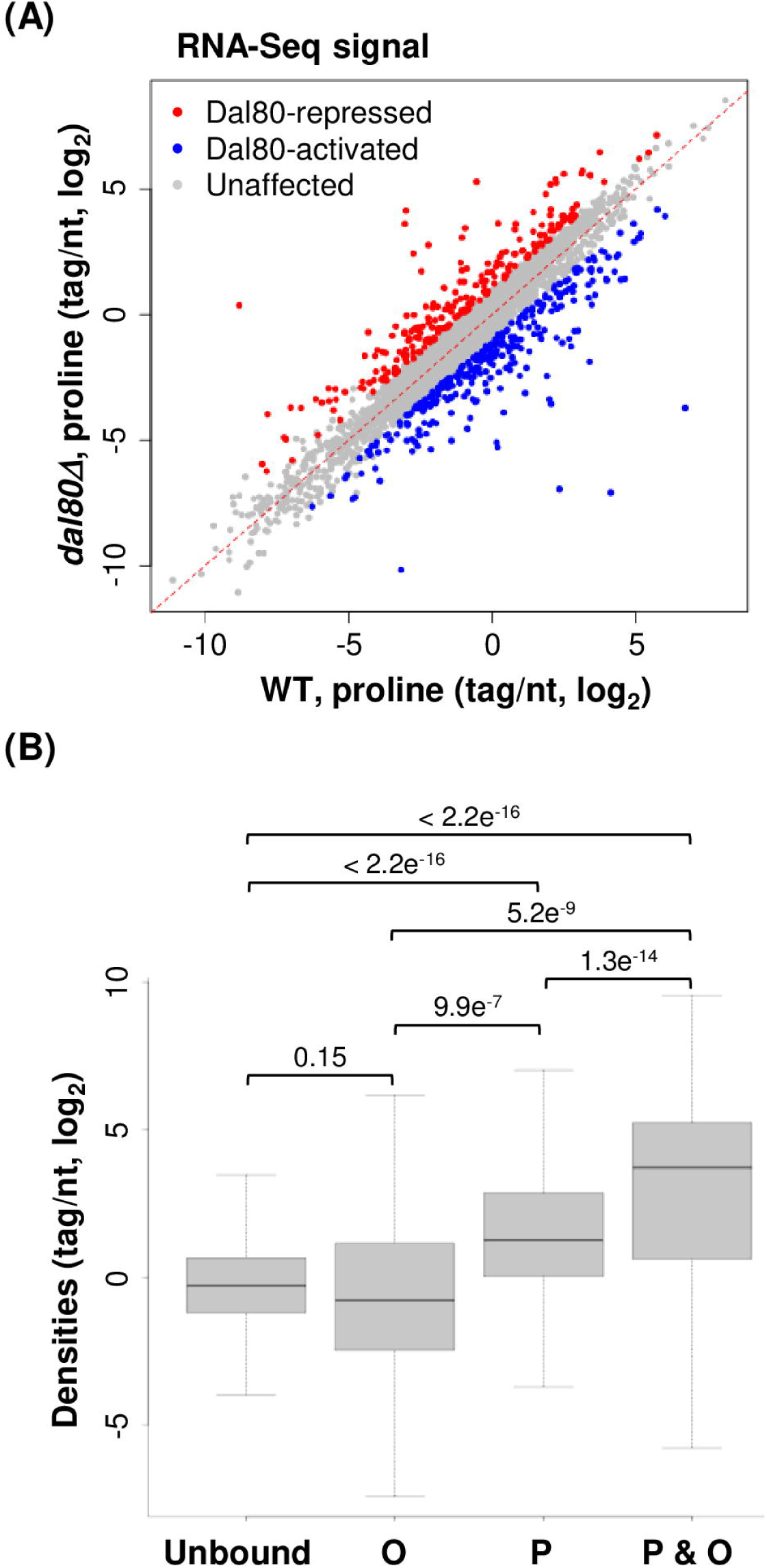
Dal80 binding across gene bodies correlates with Dal80-sensitive, high expression levels. (A) Scatter plot of densities (tag/nt, log_2_ scale) for genes in WT and *dal80*Δ cells grown in proline-containing medium. Total RNA was extracted from exponentially growing biological replicates of 25T0b (WT) and FV080 (*dal80*Δ). After rRNA depletion, strand-specific RNA-Seq libraries were constructed and then sequenced. Tag densities were computed using uniquely mapped reads. Dal80-regulated genes were identified on the basis of a min. 2-fold change (up or down) in the *dal80*Δ mutant relative to the WT control, with a *P* <0.05 upon differential expression analysis using DESeq [75]. Unaffected (n=5252), Dal80-repressed (n=232) and Dal80-activated (n=314) genes are shown as grey, red and blue dots, respectively. (B) Box-plot of densities (tag/nt, log_2_ scale) in the WT strain grown in proline-containing medium, for genes of the unbound, “O”, “P” and “P & O” classes. *P*-values obtained upon Wilcoxon rank-sum test are indicated.

Analysis of mRNA levels in proline-grown cells revealed a significant correlation between Dal80 binding and regulation at the mRNA level: 39.6% (216/546) of Dal80-regulated genes were bound by Dal80 at their promoter and/or across the ORF, which is significantly more than expected if Dal80 binding and regulation were independent (*P*<0.00001, Chi-square test; S2 Fig). More importantly, additional differences are emerging between the different classes of genes. First, gene expression was found to be globally similar between the unbound genes and those of the “O” class (*P*=0.15, Wilcoxon rank-sum test; Fig 3B). Second, Dal80 binding at the promoter of genes correlated with significantly increased expression levels, in comparison to the unbound genes (*P* < 2.2e^-16^, Wilcoxon rank-sum test; Fig 3B). Third, the genes bound by Dal80 both at the promoter and across the ORF (“P & O” class) were even more expressed than the genes showing promoter binding only (*P* = 1.3e^-14^, Wilcoxon rank-sum test; Fig 3B). Together with the observation that genes of the “P & O” class globally showed higher Dal80-Myc^13^ ChIP-Seq signal at the promoter than those of the “P” class (Fig 2C–D), our results indicate that Dal80 spreading across gene bodies correlates with a strong recruitment at the promoter, higher sensitivity to Dal80 and high expression in proline-containing medium. Consistently, metagene analysis of Dal80-regulated genes is very similar to that of “P & O” class (S2 Fig). Noticeably, 15% (170) and 2% (75) of the genes belonging to the “P” and “unbound” classes respectively display expression levels higher than median levels of expression of the genes belonging to the “P&O” class (S2 Fig). These proportions raise even further when the first quartile (25%) of the “P&O” class is considered (S2 Fig), suggesting that elevated expression levels does not automatically imply binding at ORF.

In conclusion, our genome-wide analyses revealed a significant correlation between Dal80 recruitment to the chromatin and Dal80-sensitive expression. We also showed that Dal80 binding both at the promoter and across the gene body correlates with high levels of gene expression.

### Dal80 binding across the ORF of well-characterized NCR-sensitive genes

In order to validate our genome-wide observations at the level of known NCR-sensitive genes, we characterized the binding profiles of Dal80-Myc^13^ along four NCR-sensitive genes of the “P & O” class (see snapshots of ChIP-Seq signals in Fig 4A–D). We chose the NAD^+^- dependent glutamate dehydrogenase gene (*GDH2*) and three permease-coding genes (*GAP1, DAL5* and *MEP2*), aware that their regulation could differ [37,39,47] and that GATA sites are differently represented throughout these genes (Fig 4A–D).

**Fig 4.**
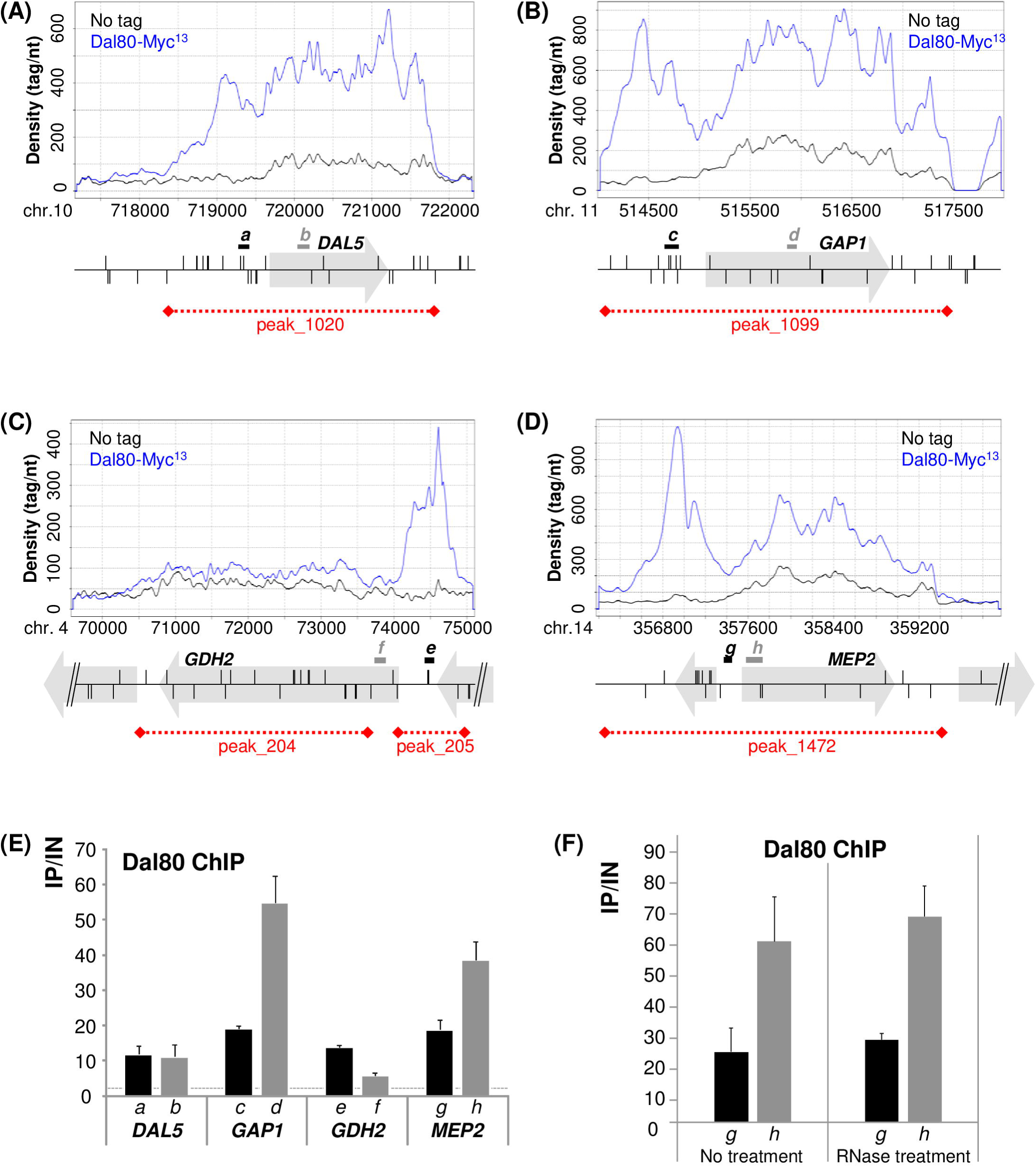
Dal80 binding across the ORF of well-characterized NCR-sensitive genes. (A-D) Position of GATA sites (vertical segments above -sense GATA sites- or below-antisense GATA sites- the *locus* line) and qPCR probes (black and grey lettered segments) along 4 NCR-sensitive loci: *DAL5* (A), *GAP1* (B), *GDH2* (C) and *MEP2* (D). (E) Cells of FV078 (wt *DAL80-MYC^13^*) strain were grown in proline as unique nitrogen source. Anti-Myc ChIP was performed as described in Experimental procedures. Each histogram represents the average (x10 000) of the value IP/IN (input or total chromatin) of at least 3 independent cultures on which two IPs were performed. The associated error bars correspond to the standard error (DAL5_P1-P2_, n=6; DAL5_O3-O4_, n=5; GAP1_P1-P2_, n=4; GAP1_O9-O10_, n=4; GDH2_P1-_P2, n=4; GDH2_O5-O6_, n=4; MEP2_P9-P10_, n=11; MEP2_O1-O2_, n=7). Black histograms represent qPCR performed on promoter regions whereas grey ones indicate qPCR data obtained within genes. Dotted line represents the highest no tag control levels. Primers are described in S10 Table: a, DAL5_P1-P2_; b, DAL5_O3-O4_; c, GAP1_P1-P2_; d, GAP1_O9-O10_; e, GDH2_P1-P2_; f, GDH2_O5-O6_; g, MEP2_P9-P10_; h, MEP2_O1-O2_. (F) ChIP analysis was conducted as in Panel E, with or without RNase treatment before immunoprecipitation (n=3).

As expected from our genome-wide observations, Dal80 bound not only the promoter, but also the coding region of *DAL5, GAP1, GDH2* and *MEP2* in proline-grown cells (Fig 4E and S3 Fig). Binding of Dal80 to the *GDH2* ORF was reduced (Fig 4E), which is consistent with the ChIP-Seq profile (Fig 4D). We ruled out any erroneous interpretation of our data upon showing that, using the same experimental design, the Gal4-Myc^13^ protein only occupied the *GAL10* promoter, and not the coding region (S3 Fig). Moreover, intragenic GATA factor binding did not rely on an intermediate RNA molecule, since the Dal80-Myc^13^ signal across the *MEP2* gene body was unaffected upon RNAse treatment of the chromatin extracts before the immunoprecipitation (Fig 4F and S3 Fig).

The results of the ChIP-qPCR show that the level of Dal80 binding also varied from one gene to another. As striking example, Dal80 was more present at the *GAP1* and *MEP2* ORFs, which are more highly expressed than *DAL5* in proline-grown cells (S6 Table). This is consistent with the correlation observed at the genomic level between Dal80 binding both to the promoter and across gene bodies and high expression levels (Fig 3B). Altogether, these observations prompted an important mechanistic question: how can Dal80 be localized to gene bodies?

#### Dal80 binding within gene bodies occurs independently of intragenic GATA sites

Like most NCR-sensitive genes, the four genes described above are all characterized by stretches of GATA sequences in their promoters (Fig 4A–D). A possible explanation for Dal80 binding across gene bodies could be that functional, intragenic GATA sequences, directly recruit Dal80. Given the low complexity of the GATA consensus, the probability that a gene is devoid of intragenic GATA site is very low. Indeed, only 10% of all yeast ORFs have no GATA site, and most genes contain an average of 5 intragenic GATA sites (S3 Fig). However, according to current models, GATA sites should be present in stretches of at least two in order to be functional *in vivo*, which is not the case within *DAL5, GDH2, GAP1* and *MEP2* ORFs, that contain some GATA sites, but relatively distant from each other. Interestingly, 23 of the 144 P&-O-bound genes did not contain any GATA site in their ORF, hence showing that Dal80 spreading across the body of genes does not necessarly require intragenic GATA sites, at least for these genes.

In order to test if the presence of a NCR-sensitive promoter could confer intragenic Dal80 binding across the body of a non-NCR-sensitive gene, devoid of any GATA cluster, we placed the *URA3* ORF under the control of the *MEP2* promoter, at the *MEP2 locus* (P*_MEP2_*- *URA3*; Fig 5A). Only two GATA sites are present in the *URA3* ORF, and separated by 300bp. In this strain, the expression of *URA3* became NCR-sensitive and followed wild type *MEP2* expression (S4 Fig), correlating with PolII recruitment over the *URA3* ORF (S4 Fig). In these conditions, we observed Dal80-Myc^13^ binding at the promoter of *MEP2* and also across *URA3* (Fig 5B). Importantly, Dal80 was not detected across *URA3* when it was expressed from its native *locus*, under the control of its promoter (Fig 5B). These results show that significant recruitment of Dal80 at an ORF region must be preceded by promoter recruitment.

**Fig 5.**
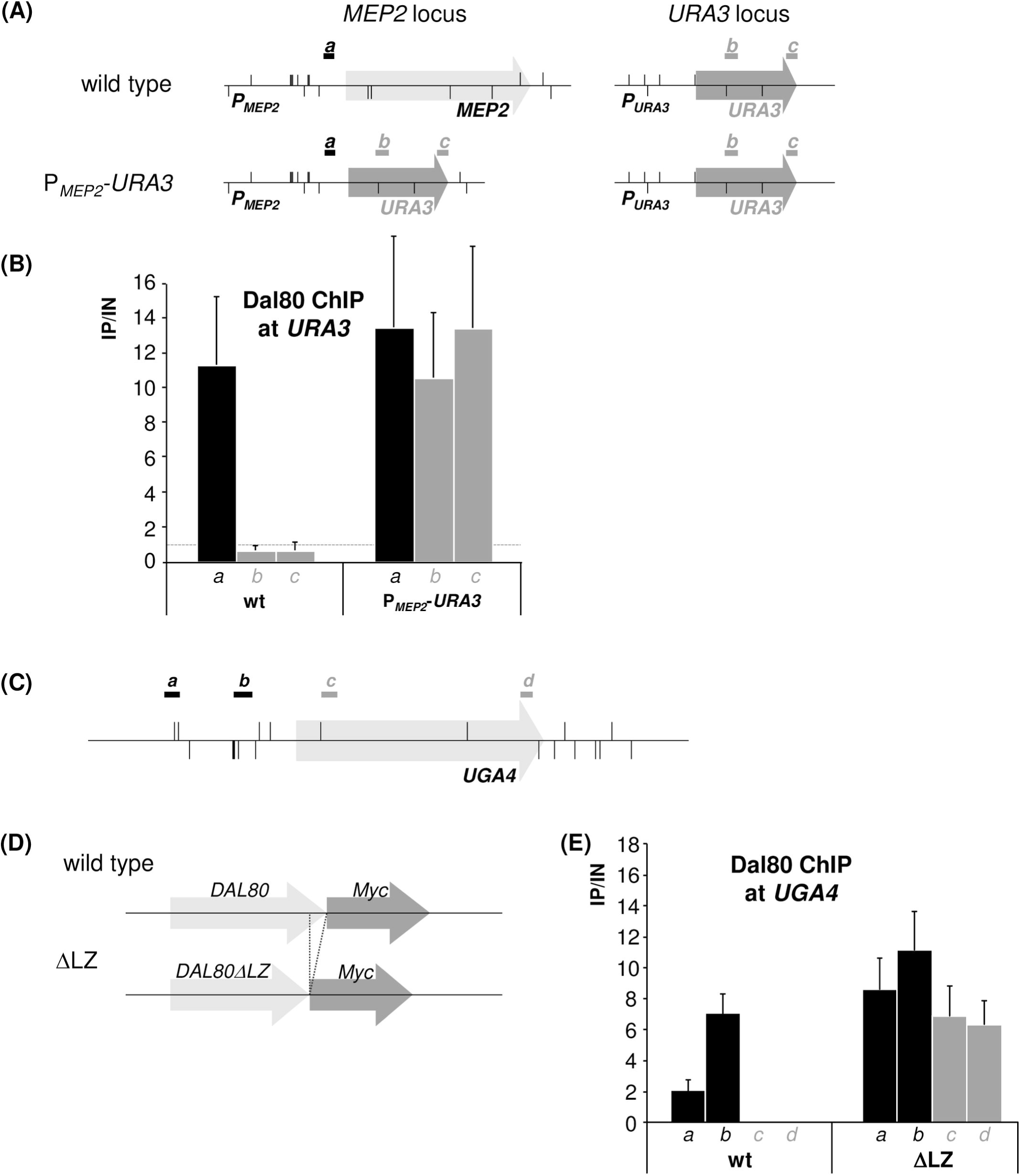
Dal80 binding within gene bodies occurs independently of intragenic GATA sites. (A) Schematic representation of the P*_MEP2_-URA3 locus*, GATA sites (vertical segments above -sense GATA sites- or below -antisense GATA sites- the *locus* line) and qPCR probe positions (black segments, *MEP2* promoter; grey segments, *URA3* ORF). (B) Occupancy of Dal80-Myc^13^ at the *MEP2* promoter and *URA3* ORF at the *MEP2* and *URA3 loci* in wild type and P*_MEP2_*-*URA3* strains. Cells of *DAL80-MYC^13^* wild type (FV078) and P*_MEP2_*-*URA3* (FV808) strains were cultivated at 29°C in the presence of proline (Pro) as unique nitrogen source. ChIP analysis was conducted as in Fig 4. The first histogram summarizes the data obtained by qPCR probing the promoter of *MEP2* (a, MEP2_P9-P10_), whereas the two following histograms are generated by qPCR performed with oligos positioned in the ORF of *URA3* (b, URA3_O1-O2_; c, URA3_O3-O4_), either at the *URA3 locus* (in wild type strain), or both at the *URA3* and *MEP2 loci*. Dotted line represents the highest no tag control levels. The associated error bars correspond to the standard error (n=6). (C) Schematic representation of the *UGA4 locus*, GATA sites (vertical segments above -sense GATA sites- or below -antisense GATA sites- the *locus* line) and qPCR probe positions (black segments, promoter; grey segments, ORF). (D) Wild type and truncated (ΔLZ) versions of Dal80-Myc^13^ used for the ChIP experiment at the *UGA4 locus*. (E) Occupancy of the *UGA4 locus* by Dal80-Myc^13^. Cells of FV078 (*DAL80-MYC^13^*; wt) and FV136 (*DAL80*Δ*LZ-MYC^13^*; ΔLZ) strains were grown at 29°C in the presence of proline (Pro) as unique nitrogen source. ChIP analysis was conducted as described in Fig 4. Black histograms represent qPCR performed with promoter-specific primers (a, UGA4_P1-P2_; b, UGA4_P9-P10_) whereas grey ones indicate qPCR data obtained with ORF-specific primers (c, UGA4_O1-O2_; d, UGA4_O3-O4_) at the *UGA4 locus*. The associated error bars correspond to the standard error (n=6).

Finally, we analyzed Dal80 binding at the *UGA4 locus*, another well-characterized NCR-sensitive gene, which is devoid of intragenic GATA site clusters (Fig 5C). *UGA4* expression is induced by GABA (γ-aminobutyric acid) and is strongly repressed by Dal80 in the absence of the inducer [48]. Therefore, to derepress *UGA4* without inducer and to analyze Dal80 binding along *UGA4*, we deleted the repressive C-terminal leucine zipper domain of Dal80 (Dal80ΔLZ-Myc^13^), known for its homodimerization and heterodimerization with the Gzf3 transcriptional repressor [17,27](Fig 5D). As a control experiment, *UGA4* expression was quantified in Dal80-Myc^13^ and Dal80ΔLZ-Myc^13^ cells. As expected, *UGA4* mRNA level was very low in Dal80-Myc^13^ cells, whereas deletion of Dal80 leucine zipper led to a strong derepression of *UGA4* (S4 Fig). Analysis of PolII occupancy across *UGA4* using ChIP confirmed that elevated mRNA levels correlate with increased transcription (S4 Fig). In these conditions, full-length Dal80-Myc^13^ binding was restricted to the *UGA4* promoter, while Dal80ΔLZ-Myc^13^ binding was detected at the promoter and across the body of *UGA4* (Fig 5E), correlating with the elevated transcript and PolII levels. Surprisingly, the leucine zipper of Dal80 and consequently, its dimerization, needed for *UGA4* repression, were not required for its spreading across *UGA4* gene body. Importantly, these results show that promoter binding is not sufficient to confer intragenic binding, but suggest that transcription activation is required.

Together, these data indicate that Dal80 binding at the promoter and across the body of NCR-sensitive genes correlated with expression levels and required transcription activation.

### Transcription elongation is necessary for Dal80 binding across *MEP2*

We hypothesized that the elongating PolII complex could be responsible for Dal80 spreading beyond Dal80-bound promoters.

To assess whether the transcriptionally active PolII complex is necessary for Dal80 binding across the *MEP2* gene, we used an *rpb1-1* strain, known to rapidly involve a cessation of PolII activity when cells are shifted to a restrictive temperature of 37°C [49,50]. We analyzed Dal80-Myc^13^ binding along *MEP2* in WT and *rpb1-1* cells. When *rpb1-1* cells were shifted at 37°C for 1hr, *MEP2* mRNA and PolII levels showed a 2-fold (S5 Fig) and >10-fold decrease (S5 Fig), respectively, reflecting the expected elongation defect when *rpb1-1* cells are shifted in non-permissive conditions. In the same conditions, we observed a significant >5-fold reduction of Dal80-Myc^13^ levels across the *MEP2* ORF, while the binding at the promoter was not affected (Fig 6A–B). This result suggests that Dal80 spreading across the body of NCR-sensitive genes would be a consequence of the PolII-driven elongation process.

**Fig 6.**
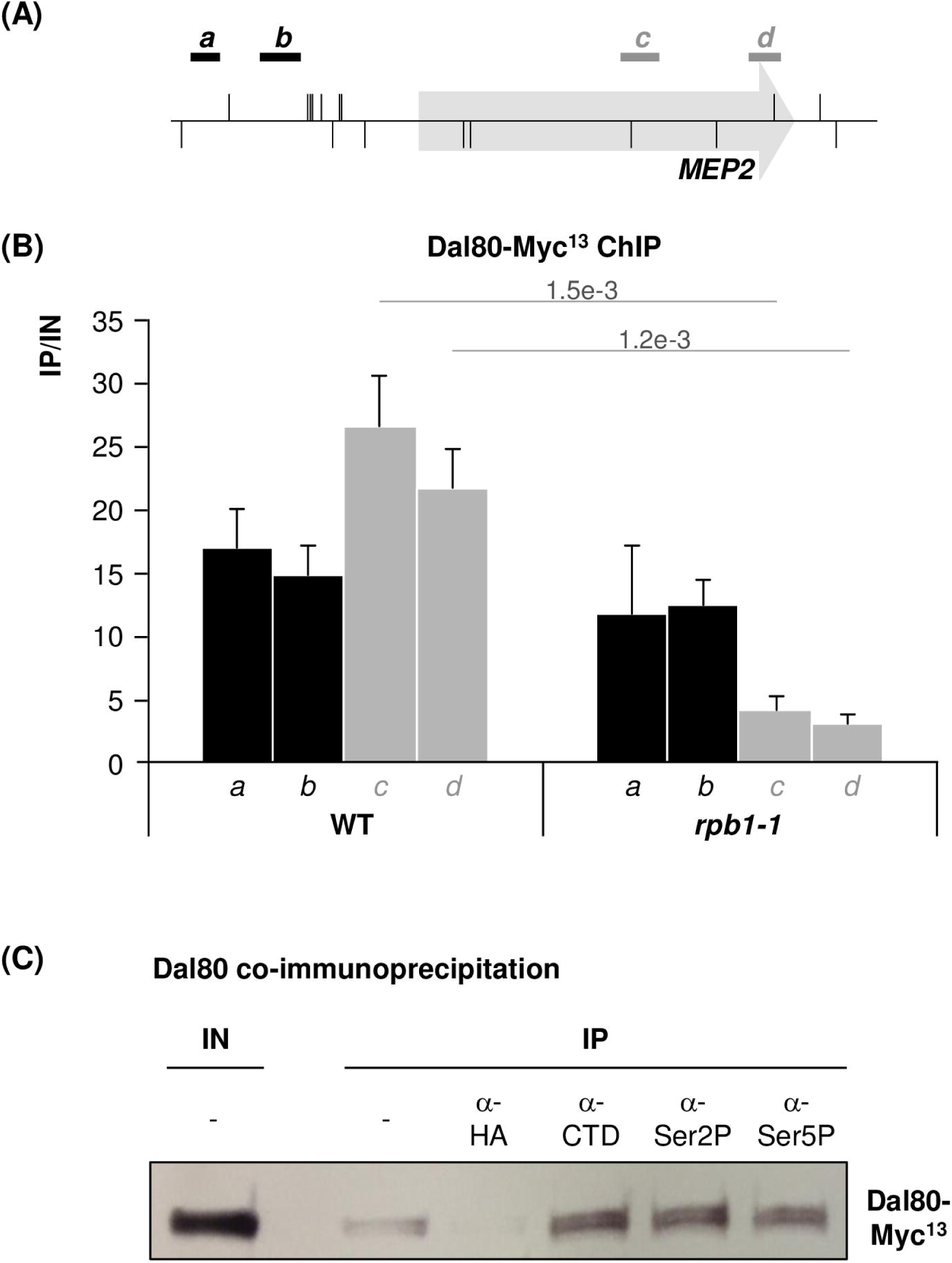
Transcription elongation is necessary for Dal80 binding across *MEP2*. (A) Schematic representation of the *MEP2 locus*, GATA sites (vertical segments above -sense GATA sites- or below -antisense GATA sites- the *locus* line), and qPCR probe positions (black segments, promoter; grey segments, ORF). (B) Occupancy of Dal80-Myc^13^ at the *MEP2 locus* when transcription elongation is impaired. Wild type (FV673) or *rpb1-1* (FV675) *DAL80-MYC^13^* cells were grown at 29°C in the presence of proline (Pro) as unique nitrogen source, and shifted at 37°C for one hour at mid-log phase. ChIP analysis was conducted as in Fig 4. The first 2 black columns represent qPCR with 2 primer pairs within the *MEP2* promoter (a, MEP2_P5-P6_; b, MEP2_P13-P14_) and the 2 following grey ones correspond to qPCR data obtained with 2 primer pairs within the *MEP2* ORF (c, MEP2_O11-O12_ d, MEP2_O9-O10_). The associated error bars correspond to the standard error (n=6). (C) Coimmunoprecipitation of Dal80-Myc^13^ with different phosphoforms of PolII. Total proteins were extracted, immunoprecipitated with the indicated antibodies, and subjected to anti-Myc western blot analysis.

To get insights into the mechanism by which Dal80 associates to gene bodies, we tested whether it physically interacts with the transcriptionnaly engaged form of PolII (Fig 6C). Total protein extracts from Dal80-Myc^13^ cells were immunoprecipitated with antibodies directed against the PolII CTD and its phospho-forms Ser2P and Ser5P, respectively characteristic of elongating and initiating PolII forms. All three antibodies enabled effective immunoprecipitation, whereas no antibody and nonspecific antibody controls generated a lower or no signal at all. Thus, Dal80 physically interacts with phosphoforms of the PolIII, suggesting a strong association with PolII engaged in active transcription from initiating to elongating polymerase.

Together, our data indicate that Dal80 spreading across the body of NCR-sensitive genes depends on active transcription and that Dal80 interacts with the transcriptioanlly active form of PolII, supporting a model where Dal80 spreading across the body of highly expressed, NCR-sensitive genes might be the result of Dal80-PolII association at post-initiation transcription phases.

## Discussion

Eukaryotic GATA factors belong to a small but important family of DNA binding proteins involved in development and response to environmental changes in multicellular organisms in unicellular organisms, respectively. In yeast, four GATA factors are involved in Nitrogen Catabolite Repression (NCR), controlling gene expression in response to nitrogen source availability. One of them, the Dal80 repressor, itself NCR-sensitive, acts to modulate the intensity of NCR responses.

Over the past decade, a number of studies have screened the genome aiming at gathering an inventory of genes regulated by the nitrogen source. Although >500 genes have been shown to be differentially expressed upon change of the nitrogen source [39,46], the list of NCR-sensitive genes was reduced to about 100, based on their sensitivity to GATA factors [37,39,42,45], suggesting that the number of Dal80 targets would be situated in that range. Here, our ChIP-Seq analysis identified more than 1200 Dal80-bound promoters, which considerably extends the list of potential Dal80 targets. In fact, the number of Dal80-bound promoters could even have been more important. Indeed, the GATA consensus binding site is rather simple and short, so that in yeast, a total number of 10,000 putative binding sites can be found in all protein-coding gene promoters, 2930 promoters having at least 2 GATA sites, which is thought to be a prerequisite for *in vivo* binding and function of the GATA factors. The difference between the number of promoters with ≥2 GATA sites and the number of Dal80-bound promoters suggests the existence of a selectivity for Dal80 recruitment. This selectivity could rely on promoter architecture and/or chromatin structure, conditioning the requirement for auxiliary DNA binding factors that would stabilize Dal80 at some promoters. Moreover, although we observed a significant correlation between Dal80 binding and regulation, the expression of most of the Dal80-bound genes was not affected in a *dal80*Δ mutant strain. Again, Dal80-dependence for transcribing these genes, as well as their NCR sensitivity, could require the presence of cofactors which are not produced or inactive under the tested growth conditions. In mammals, GATA factors also display an extraordinary complexity in the relationships between binding and expression regulation. Like Dal80, GATA-1 and GATA-2 only occupy a small subset of their abundant binding motif throughout the genome, and the presence of the conserved binding site is insufficient to cause GATA-dependent regulation in most instances [51]. GATA-1 binding kinetics, stoichiometry and heterogenous complex formations, conditioned by composite promoter architecture, influence its transcriptional activity and hence diversify gene expression profiles [51].

Given the high conservation at the amino acid level between the DNA binding domain of the four yeast NCR GATA factors, it is likely that they all recognize identical sequences (GATAA, GATAAG or GATTAG). This consensus has been largely validated in the past using gene reporter experiments, mutational analyses and *in vitro* binding experiments on naked DNA. Nonetheless, of the 1269 bound promoters, 48% contained at least two GATA sites, a proportion that is not different from that observed among unbound promoters, and the amount of GATA sites per promoter was not different between the two groups either. In addition, Dal80 recruitment was found to occur independently of the presence of GATA sites in 20% of Dal80-bound promoters, as also previously observed in mammalian cells [10,52]. Future experiments will be required to decipher the molecular mechanism by which Dal80 can be recruited to these GATA-less promoters.

Unexpectedly, although Dal80 has always been described as a repressor, we identified 314 genes that are positively regulated by Dal80 (their expression is significantly decreased upon Dal80 deletion). These genes are distinct from the NCR-sensitive genes: only 4 belong to the NCR-sensitive list of [39] and are significantly enriched in amino acid biosynthetic processes, resembling the amino acid starvation response mediated by the Gcn4 transcriptional activator. Interestingly, the promoter of 122/314 Dal80-activated genes contain Gcn4-binding sites (S3 Fig), and this group of 314 Dal80-activated genes is significantly enriched for genes regulated by the General Amino Acid Control (GAAC; YeastMine Gene List, Publication Enrichment, p<1.6e-13), through the Gcn4 activator. Interconnections between NCR and GAAC have already been demonstrated, mostly at the level of nitrogen catabolism control: 1-a large number of non-preferential nitrogen sources leads to increased transcription of GAAC targets [39]; and 2- Gcn4 contributes, with Gln3, to the expression of NCR-sensitive genes [53,54]. However, this is the first time that evidence are provided suggesting a positive function for the negative GATA factor at the level biosynthetic gene expression.

The most striking and unexpected finding of this work is the observation that for 189 genes, Dal80 occupied the gene body (“O” and “P & O” classes). The 45 genes displaying Dal80 intragenic occupancy with no recruitment at the promoter (“O” class) were globally neither Dal80-sensitive nor NCR-sensitive, similar to the unbound genes. Furthermore, we also noted that a substantial fraction of these genes correspond to small dubious ORFs, near or even overlapping the promoter, Dal80-bound, of the adjacent gene. In these cases, the limited resolution of the ChIP-Seq technique, combined to the small size of these genes, might have allowed them to pass through the filters of the procedure to identify the genes showing Dal80 intragenic binding. Overall, these observations suggest that the existence of the “O” class is likely to be physiologically irrelevant. Dal80 binding at the promoter and spreading sacross the body of the 144 genes belonging to the “P & O” class correlated with high expression levels and sensitivity to Dal80-dependent repression. The observation that a considerable fraction of “P” class genes have similar or higher expression levels than most of the “P&O” class genes suggest that the well-known artifact of hyper-chippable regions is not applicable here [55]. Our observations were also experimentally confirmed using ChIP experiments, at the level of well-characterized NCR-sensitive genes. Our data indicate that Dal80 spreading across genes bodies does not rely on intragenic GATA sites. Rather, promoter binding appears to be required but not sufficient: Dal80 spreading across genes depends on active transcription. Indeed, the observation that impaired PolII elongation correlates with decreased intragenic binding further indicates that Dal80 binding across gene bodies depends on ongoing transcription elongation. Consistently, we detected a physical interaction between Dal80 and transcriptionnaly active forms of PolII. Together, our data leads us to propose a model where Dal80 could travel from the promoter of highly expressed, NCR-sensitive genes through the gene body by accompanying the elongating PolII complex (Fig 7). Additional investigations will be required to define which domain of Dal80 is responsible for the interaction with the transcription machinery, to determine whether there is any causal relationship between Dal80 intragenic binding and high expression levels, and to decipher the potential role of Dal80 during transcription elongation.

**Fig 7.**
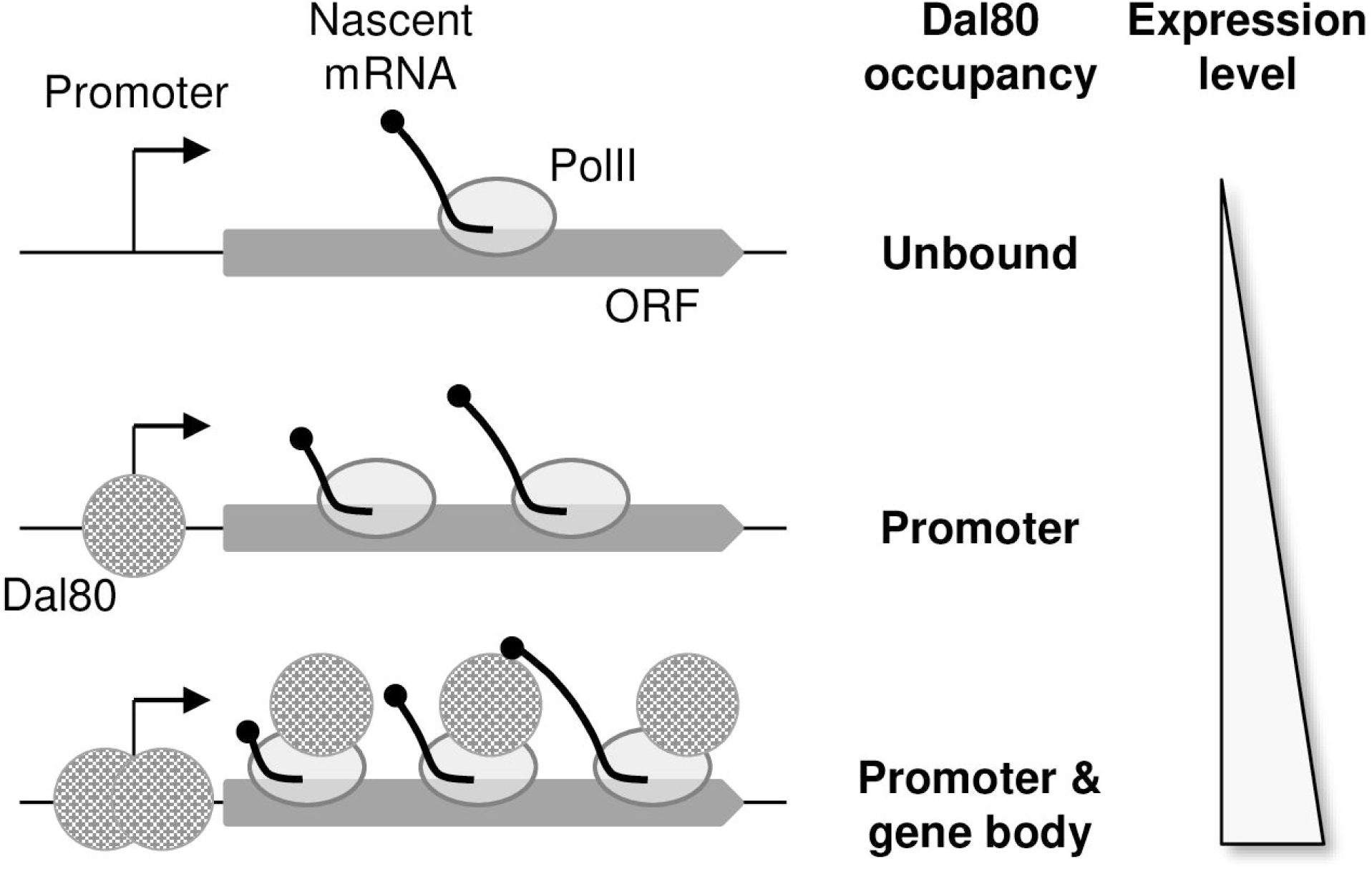
Model summarizing the correlations observed between Dal80 binding, Dal80 dependency and expression levels.

Whereas the binding of elongation factors across gene bodies has been thoroughly documented [56], it has also been described for few specific transcription factors. For example, Gal4 was reported to bind to its consensus DNA target within the *ACC1* ORF, but the authors concluded that the observed transcriptional repression of the *ACC1* gene was most likely resulting from random *GAL4* binding “noise” over the genome, thus having no physiological explanation for this ORF-bound transcription factor [57]. Likewise, Gcn4 was detected across the *PHO8* ORF, with concomitant recruitment of the SAGA complex, but without any impact on gene expression [58]. More recently, binding of the Gcn4 transcription factor to its consensus site at some ORFs, when located in proximity of the transcriptional start site, was found to play a consistent role in controlling embedded cryptic promoters in yeast, thereby affecting Gcn4-dependent transcription of some genes [59].

A recent study has identified CTD phosphorylation of polII as a hub that optimizes transcriptome changes to adequately balance optimal growth and stress tolerance responses [60]. The addition of nitrogen to nitrogen-limited cells rapidly results in the transient overproduction of transcripts required for protein translation (stimulated growth) whereas accelerated mRNA degradation favours rapid clearing of the most abundant transcripts, like those involved in high affinity permease production, that are highly expressed NCR-sensitive genes, for example [46]. The involvement of the Nrd1-Nab3-Sen1 (NNS) and TRAMP complexes in these regulatory responses has been envisioned very recently [61,62]; deadenylation, decapping and exonuclease mutants display impaired *GAP1* mRNA clearance upon nitrogen upshift [63]. Thus, a possible role of Dal80 (and possibly of the other GATA factors) binding along highly expressed genes could be to transmit nutritional signals to the RNA decay machinery for rapid and efficient remodeling of gene expression in response to changing nitrogen supplies. Here, an attractive model (Fig 7) would hypothesize that Dal80 binding to polII could prevent the decay of the dedicated stress-response mRNAs to enhance their cellular life-cycle by regulating/repressing RNA decay factors that might associate otherwise on the polymerase. Other elongation-related processes, like histone modification and chromatin remodeling [64,65], mRNA export [66] or roadblock termination [67] can also be proposed.

In this respect, in human cells, GATA factors are also reported to occupy non-canonical sites within the genome, further reinforcing that they can be recruited to the chromatin independently of their motif [10,52]. In addition, 43% of the GATA1 peaks were collected among exon, introns and 3’UTR of coding genes in human erythroleukemia cells [52]. It is tempting to hypothesize that GATA factors could have a dual or synergistic role during transcription, i.e. recruiting/stabilizing the PIC complex as for any classical transcription factor in the promoter/enhancer regions and promoting elongation at a post initiation step interacting with the RNAPII. Interestingly, GATA1 in erythropoid cells interacts with the mediator [68], a complex also known to travel with PolII [69].

## Materials and methods

### Experimental model and subject details

Experiments were conducted using *S. cerevisiae* strains of the FY genetic background. The strains used are listed in S9 Table. Dal80 was tagged with 13 copies of the *c-myc* epitope (Myc^13^) as described [70] using primers listed in S9 and S10 Tables. The *P_MEP2_-URA3* allele in strains FV806-808 was created by amplification of the *URA3* gene using the same strategy, with primers listed in S9 and S10 Tables.

Cultures were grown at 29°C to mid-log phase (A_660nm_ = 0.5) in YNB (without amino acids or ammonia) minimal medium containing the indicated nitrogen source at a 0.1% final concentration, glucose (3%) and the appropriate supplements (20 µg/ml uracil, histidine and tryptophan) to cover auxotrophic requirements.

### Chromatin immunoprecipitation

Cell extracts and immunoprecipitations were conducted as described [23] using primers listed in S10 Table. The cells (100-ml cultures grown to an absorbance (A660 nm = 0.6) corresponding to 6 × 106 cells/ml) were treated with 1% formaldehyde for 30 min at 25 °C and mixed by orbital shaking. Glycine was then added to a final concentration of 500 mm and incubation continued for 5 min. The cells were collected, washed once with cold 10 mm Tris- HCl, pH 8, washed once with cold FA-SDS buffer (50 mm HEPES-KOH, pH 7.5, 150 mm NaCl, 1 mm EDTA, 1% Triton X-100, 0.1% sodium deoxycholate, 0.1% SDS, 1 mm phenylmethylsulfonyl fluoride), and resuspended in 1 ml of cold FA-SDS buffer. An equal volume of glass beads (0.5 mm in diameter) was added, and the cells were disrupted by vortexing for 30 min in a cold room. The lysate was diluted into 4 ml of FA-SDS buffer, and the glass beads were discarded. The cross-linked chromatin was then pelleted by centrifugation (17,000 × g for 35 min), washed for 60 min with FA-SDS buffer, resuspended in 1.6 ml of FA-SDS buffer for 15 min at 4 °C, and sonicated three times for 30 s. each (Bioruptor, Diagenode) to yield an average DNA fragment size of 700 base pairs. Finally, the sample was clarified by centrifugation at 14,000 × g for 30 min and diluted 4-fold in FA-SDS buffer, and aliquots of the resultant chromatin containing solution were stored at –80 °C. Myc^13^-tagged proteins were immunoprecipitated by incubating 100 μl of the chromatin containing solution for 180 min at 4°C with 2 μl of mouse anti-Myc antibodies (SC-40, Santa Cruz) prebound to 10 μl of Dynabeads Pan Mouse IgG (Dynal) according to the manufacturer’s instructions. Immune complexes were washed six times in FA-SDS buffer and recovered by treating with 50 μl of Pronase Buffer (25 mm Tris, pH 7.5, 5 mm EDTA, 0.5% SDS) at 65 °C with agitation. Input (IN) and immunoprecipitated (IP) fractions were then subjected to Pronase treatment (0.5 mg/ml; Roche Applied Science) for 60 min at 37°C, and formaldehyde cross-links were reversed by incubating the eluates overnight at 65°C. Finally, the samples were treated with RNase (50 μg/ml) for 60 min at 37°C. DNA from the IP fractions was purified using the High Pure PCR Product Purification Kit (Roche Applied Science) and eluted in 50 μl of 20 mm Tris buffer, pH 8. IN fractions were boiled 10 min and diluted 500-fold with no further purification prior to quantitative PCR analysis.

### Quantitative RT-PCR

Quantitative RT-PCR was performed as described previously [23] using primers listed in S10 Table. Total RNA was extracted from 4-ml cultures and cDNA was generated from 100 to 500 ng of total RNA using a RevertAid H Minus first-strand cDNA synthesis kit with oligo(dT)_18_ primers from Fermentas using the manufacturer’s recommended protocol. cDNAs were subsequently quantified by RT-PCR using the Maxima SYBR green qPCR master mix from Fermentas.

### Co-immunoprecipitation

Cultures (100 ml) were harvested, washed once in 50 mM Tris, pH 8, and resuspended in 1ml of buffer (50 mM Tris, pH 8, 150 mM NaCl, 5 mM EDTA, 0.05% NP-40, 1 mM phenylmetyhlsulfonyl fluoride, and complete protease inhibitor cocktail tablets [Roche]). Lysis was performed by shaking with 425–600 μm acid-washed glass beads (Sigma) on an IKA Vibrax VXR orbital shaker at maximum speed for 30 min at 4°C. Cell debris and glass beads were removed by centrifugation. Immunoprecipitation was performed by incubating 200 μl of total cell extracts with 20 μl of Dynabeads PAN mouse immunoglobulin G (Invitrogen) that were preinucbated with anti-HA (SCBT, SC-7392), anti-CTD (SCBT, CTD4H8), anti-Ser2P (BioLegend, H5) or anti-Ser5P (BioLegend, H14) antibodies and 20 μl of 1% phosphate-buffered saline-bovine serum albumin for 2 h under orbital shaking (800 rpm) at 30°C. Immune complexes were washed three times in lysis buffer, eluted by boiling in sodium dodecyl sulfate (SDS) sample buffer, and loaded on SDS-polyacrylamide gel for anti-Myc Western blotting.

### ChIP-Seq analysis and peak-calling

ChIP-Seq analysis was performed from two biological replicates of proline-grown 25T0b (no tag) and FV078 (Dal80-Myc^13^) cells. For each condition, libraries were prepared from 10 ng of “input” or “IP” DNA using the TruSeq ChIP Sample Preparation Kit (Illumina). Single-read sequencing (50 nt) of the libraries was performed on a HiSeq 2500 sequencer.

Reads were uniquely mapped to the *S. cerevisiae* S288 C reference genome using Bowtie2 v2.1.0 [71], with a tolerance of 1 mismatch in seed alignment. Tags densities were normalized on the total number of uniquely reads mapped.

Dal80-bound regions were identified through a peak-calling procedure using version 2.0.9 of MACS [72], with a minimum false discovery rate (FDR) of 0.001.

### Total RNA-Seq

For each strain and condition, total RNA was extracted from two biological replicates using standard hot phenol procedure, ethanol-precipitated, resuspended in nuclease-free H_2_O (Ambion) and quantified using a NanoDrop 2000c spectrophotometer. Ribosomal RNAs were depleted from 1 µg of total RNA using the RiboMinus^TM^ Eukaryote v2 Kit (Life Technologies). After concentration using the Ribominus^TM^ Concentration Module (Life Technologies), rRNA-depleted RNA was quantified using the Qubit RNA HS Assay kit (Life Technologies). In parallel, rRNA depletion efficiency and integrity of both total and rRNA-depleted RNA were checked by analysis in a RNA 6000 Pico chip, in a 2100 bioanalyzer (Agilent). Strand-specific total RNA-Seq libraries were prepared from 125 ng of rRNA-depleted RNA using the TruSeq^®^ Stranded Total RNA Sample Preparation Kit (Illumina), following manufacturer’s instructions. Paired-end sequencing (2 × 50 nt) of the libraries was performed on a HiSeq 2500 sequencer. Sequenced reads were mapped to the reference genome using version 2.0.6 of TopHat [73], as described [74]. Tags densities were normalized on the total number of reads uniquely mapped on ORFs. Differential expression analysis was performed using DESeq [75].

#### Quantification and Statistical Analysis

Statistical details are to be found in the figure legends. Error bars correspond to standard error. Statistical significance tests were carried out using the Student’s t test (Microsoft Excel) when indicated.

### Availability of data and materials

Sequence data can be accessed at the NCBI Gene Expression Omnibus using accession numbers GSE86307 and GSE86325. Genome browsers for visualization of processed ChIP-Seq and RNA-Seq data are accessible at http://vm-gb.curie.fr/dal80.

Further information and requests for resources and reagents should be directed to and will be fulfilled by the lead contact, Isabelle Georis. Bioinformatics and genome wide dataset requests could also be addressed to for rapid processing.

## Acknowledgements

This work has benefited from the facilities and expertise of the NGS platform of Institut Curie.

We are grateful to all members of our labs for discussions and critical reading of the manuscript. We thank F. Vierendeels for excellent technical assistance.

## Supporting information

**S1 Fig. Genome-wide identification of Dal80-bound promoters.** (A) Box-plot of the distance between the annotated TSS and ORF start site (translation initiation codon, ATG) for protein-coding genes. (B) Snapshot of ChIP-Seq signals along a GATA-less *locus* (*ALD6*). Densities (tag/nt) are shown for the untagged (black line) and Dal80-Myc^13^ (blue line) strains. ORFs and the Dal80-Myc^13^ ChIP-Seq peak are represented by grey arrows and the red dashed line, respectively. The snapshot was produced using the VING software [76].

**S1 Table. Genome-wide identification of Dal80-bound promoters.** List of 1269 gene promoters bound by Dal80.

**S2 Table. Genome-wide identification of Dal80-bound promoters.** Overlap of genes identified in our screens and in previous genome-wide expression screens.

**S3 Table. Genome-wide identification of Dal80-bound promoters.** GO term analysis of 1269 Dal80-bound promoters.

**S4 Table. Genome-wide identification of Dal80-bound promoters.** Whole genome GATA site contents of promoters and ORFs, correlated with Dal80 binding, regulation, NCR status and presence of Gcn4-binding sites.

**S5 Table. Dal80 binds the intragenic region of a subset of genes.** Inventory of gene promoters and ORFs bound by Dal80 and their intersection.

**S6 Table. Dal80 binding across gene bodies correlates with Dal80-sensitive, high expression levels.** List of 546 Dal80-regulated genes on proline.

**S7 Table. Dal80 binding across gene bodies correlates with Dal80-sensitive, high expression levels.** Gene ontology term analysis of the 232 genes repressed by Dal80 on proline.

**S8 Table. Dal80 binding across gene bodies correlates with Dal80-sensitive, high expression levels.** Gene ontology term analysis of the 314 genes activated by Dal80 on proline.

**S2 Fig. Dal80 binding across gene bodies correlates with Dal80-sensitive, high expression levels.** (A) Contingency table showing the number of Dal80-regulated or unaffected genes (defined upon RNA-Seq analysis in WT and *dal80*Δ cells) among the Dal80-bound and unbound genes (identified upon ChIP-Seq analysis in untagged and Dal80-Myc^13^ cells). Results that were experimentally observed or that were expected in case of independence are indicated in bold and in brackets, respectively. (B) Metagene view of the ChIP-Seq signal along the 216 Dal80-regulated and -bound genes (solid lines), and the 330 Dal80-regulated and -unbound genes (dashed lines), in the untagged (black) and Dal80-Myc^13^ (blue) strains. The shading surrounding each line denotes the 95% confidence interval. (C) Density-plot of RNA-Seq signal (tag/nt, log2 scale) in the WT strain grown in proline-containing, for genes of the “unbound” (blue, n=4484), “P” (red, n=1125) and P&O” (black, n=144) classes. Y-axis: proportion of genes for each class. The highlighted areas correspond to the 75 (2%) and 170 (15%) genes of the “unbound” and “P” classes, respectively, showing a signal higher than the median of the “P&O” class. A box-plot representation of the same RNA-Seq signals is shown on the top of the density-plot. (D) Same as c., highlighting the 949 (21%) and 632 (56%) genes of the “unbound” and “P” classes, respectively, showing a signal higher than the first quartile value for the “P&O” class.

**S3 Fig. Dal80 binding across the ORF of well-characterized NCR-sensitive genes – Control experiments.** (A) Cells of FV078 (wt *DAL80-MYC^13^*) strain were grown in glutamine (Gln) or proline (Pro) as unique nitrogen sources. Anti-Myc ChIP was performed as described in Experimental procedures. Each histogram represents the average (x10 000) of the value IP/IN (input or total chromatin) of at least 3 independent cultures on which two IPs were performed. The associated error bars correspond to the standard error (DAL5_P1-P2_, n=6; DAL5_O3-O4_, n=5; GAP1_P1-P2_, n=4; GAP1_O9-O10_, n=4; GDH2_P1-P2_, n=4; GDH2_O5-O6_, n=4; MEP2_P9-P10_, n=11; MEP2_O1-O2_, n=7). Black histograms represent qPCR performed on promoter regions whereas grey ones indicate qPCR data obtained within genes. Primers are described in S10 Table: DAL5_P1-P2_, DAL5_O3-O4_, GAP1_P1-P2_, GAP1_O9-O10_, GDH2_P1-P2_, GDH2_O5-_O6, MEP2_P9-P10_, MEP2_O1-O2_. (B) Gal4-Myc^13^ binding at the *GAL1-10* promoter. Schematic representation of the *GAL1-10 locus*. Position of the Gal4-binding sites (vertical segments across the *locus* line) and qPCR probes (black and grey lettered segments) are indicated. Cells of FV437 (wt *GAL4-MYC^13^*) strain were grown in YP with glucose or galactose as unique carbon sources. Anti-Myc ChIP was performed as described in Experimental procedures. Each histogram represents the average (x10 000) of the value IP/IN (input or total chromatin) of 2 independent cultures. The associated error bars correspond to the standard error (n=2). Black histograms represent qPCR performed on promoter regions whereas grey ones indicate qPCR data obtained within genes. Primers are described in S10 Table. (C) Binding of Dal80- Myc^13^ at the *MEP2 locus*, in untreated or RNase-treated cells. Schematic representation of the *MEP2 locus*. Position of the GATA sites (vertical segments above -sense GATA sites- or below -antisense GATA sites- the *locus* line) and qPCR probes (black and grey lettered segments) are indicated. Cells of untagged (25T0b) or FV078 (*DAL80-MYC^13^*; wt) strains were grown at 29°C in the presence of proline (Pro) as unique nitrogen source. ChIP analysis was conducted with or without RNase treatment before immunoprecipitation. The first 3 black columns represent qPCR with 3 primer pairs within the *MEP2* promoter MEP2_P5-P6_; MEP2_P13-_ P14; MEP2_P9-P10_) and the 3 following grey ones correspond to qPCR data obtained with 3 primer pairs within the *MEP2* ORF (MEP2_O1-O2_; MEP2_O11-O12_ MEP2_O9-O10_). Each histogram represents the average (x10 000) of the value IP/IN (input or total chromatin) of independent cultures. The associated error bars correspond to the standard error (n=3).

**S4 Fig. Dal80 binding within gene bodies occurs independently of intragenic GATA sites – Control experiments.** (A) Expression of the non-NCR-sensitive gene *URA3* under the control of P*_MEP2_*. Cells of wild type (25T0b) and P*_MEP2_*-*URA3* (FV806) strains were cultivated at 29°C in the presence of proline (Pro) as unique nitrogen source. The expression of *URA3* is normalized to the expression of the *TBP1* gene. Each column corresponds to the mean value of data obtained for at least 2 independent cultures and the error bars correspond to the standard error (n=2). (B) PolII occupancy at the *URA3* and *P_MEP2_-URA3 loci*. Cells of *DAL80- MYC^13^* wild type (FV078) and P*_MEP2_*-*URA3* (FV808) strains were cultivated at 29°C in the presence of glutamine (Gln) or proline (Pro) as unique nitrogen sources. Anti-polII (CTD4H8) ChIP was performed as described in Experimental procedures. Each histogram represents the average (x10 000) of the value IP/IN (input or total chromatin) of at least 2 independent cultures on which two IPs were performed. Histograms represent qPCR performed with URA3_O1-O2_ primers (n=2). (C) *UGA4* expression in a *DAL80*Δ*LZ* mutant strain. Cells of FV078 (*DAL80-MYC^13^*; wt) and FV136 (*DAL80*Δ*LZ-MYC^13^*; ΔLZ) strains were cultivated at 29°C in the presence of proline (Pro) as unique nitrogen source. Total RNA was isolated and *TBP1*-normalized *UGA4* mRNA levels were quantified by qRT-PCR using UGA4_O1-O2_ primers. Histograms represent the averages of at least 2 independent experiments and the associated error bars correspond to the standard error (n=3). (D) Occupancy of the *UGA4 locus* by PolII. Cells of FV078 (*DAL80-MYC^13^*; wt) and FV136 (*DAL80*Δ*LZ-MYC^13^*; ΔLZ) strains were grown at 29°C in the presence of proline (Pro) as unique nitrogen source. Anti-polII (CTD4H8) ChIP analysis was conducted as in Panel B using UGA4_O1-O2_ primers (n=2).

**S5 Fig. Transcription elongation is necessary for Dal80 binding across *MEP2* – Control experiments.** (A) Effect of PolII elongation defects on *MEP2* expression. Wild type (FV673) or *rpb1-1* (FV675) *DAL80-MYC^13^* cells were grown at 29°C in the presence of glutamine (Gln) or proline (Pro) as unique nitrogen sources, and shifted at 37°C for one hour at mid-log phase. Total RNA was isolated and *TBP1*-normalized *MEP2* mRNA levels were quantified by qRT-PCR using MEP2_O9-O10_ primers. Histograms represent the average of at least 2 independent experiments and the associated error bars correspond to the standard error (n=2). (B) PolII occupancy at the *MEP2 locus*. Wild type (FV673) or *rpb1-1* (FV675) *DAL80- MYC^13^* cells were grown at 29°C in the presence of glutamine (Gln) or proline (Pro) as unique nitrogen sources, and shifted at 37°C for one hour at mid-log phase. ChIP analysis was conducted as in S4 Fig using MEP2_O9-O10_ primers (n=4).

**S9 Table. Strains used in this study.** [50]

**S10 Table. Primers used in this study.**

